# Borzoi-informed fine mapping improves causal variant prioritization in complex trait GWAS

**DOI:** 10.1101/2025.07.09.663936

**Authors:** Divyanshi Srivastava, Anya Korsakova, Qingbo Wang, Luong Ruiz, Han Yuan, David R Kelley

## Abstract

Genome-wide association studies (GWAS) have identified thousands of trait-associated loci. Prioritizing causal variants within these loci is critical for characterizing trait biology. Statistical fine mapping identifies causal variants at trait-associated loci, but linkage disequilibrium (LD) and limited GWAS sample sizes prevent the resolution of many associations. Functionally informed approaches augment fine mapping by estimating variant prior causal probabilities based on overlap with trait-relevant functional annotations. However, functional enrichment provides only an indirect proxy for variant functional impact. Sequence-to-function models directly estimate variant effects on molecular phenotypes from underlying sequence context. Borzoi is a long-context model that predicts sequence determinants of transcription, splicing, and polyadenylation across diverse tissues and cell types. Here we present Sniff, a Borzoi-informed fine-mapping approach that integrates broad genomic functional annotations with Borzoi-predicted variant effects via PolyFun to estimate variant prior causal probabilities. Applied to 15 UK Biobank traits, Sniff identifies 9.45% additional fine-mapped variants compared to PolyFun-Baseline at posterior inclusion probability (PIP) > 0.8. Sniff-prioritized variants exhibit allele-specific activity in reporter assays and are predicted to have tissue-specific activity in trait-relevant tissues. For most traits, genes nominated by Sniff receive higher scores from the orthogonal gene prioritization method PoPS compared to genes nominated using functional annotations alone. Because differentially prioritized variants are driven by Borzoi predictions, we leverage attribution techniques to characterize sequence features underlying fine mapping and generate mechanistic hypotheses for GWAS associations.

## 2 Introduction

Genome-wide association studies (GWAS) have identified thousands of complex trait and disease-associated loci [1, 2]. However, marginal association tests implicate multiple correlated variants at each locus due to linkage disequilibrium (LD), making causal variant identification challenging [2–4]. Bayesian fine-mapping approaches such as SuSiE [5, 6], FINEMAP [4], and CARMA [7] jointly model variant effects to estimate posterior inclusion probabilities (PIPs) for prioritizing likely causal variants. Although these approaches successfully prioritize variants at trait-associated loci, they remain unable to resolve many associations due to extensive LD and limited GWAS sample sizes.

Functionally informed fine-mapping methods leverage variant annotations including conservation scores, coding effects, minor allele frequencies (MAFs), and regulatory element overlap to guide causal variant prioritization [8, 9]. PolyFun [8] uses genome-wide baseline-LF (v2.2.UKB) annotations (N=187) [10] to estimate per-variant causal probabilities, which fit naturally as priors within Bayesian fine-mapping frameworks [4, 5, 7]. However, relying solely on functional annotations for inferring variant causal probabilities has several limitations. First, variants overlapping regulatory elements may not influence element activity—only a fraction of nucleotides within enhancers drive activity [11–14]. Second, when multiple GWAS-associated variants lie in functionally relevant regions, annotation-based fine mapping cannot discriminate among potential causal variants. Third, variant activity in sequence motifs is context-dependent, which standard baseline annotations typically do not capture [15–19]. Fourth, regulatory annotations based on the reference genome miss gain-of-function variants outside annotated functional regions [16, 20].

DNA sequence-based deep neural networks can directly estimate variant functional consequences rather than relying on annotation overlap as a proxy [21–30]. These models are trained to predict diverse molecular phenotypes including chromatin accessibility [23, 27], transcription factor binding [21], splicing [22, 28], transcription initiation [31, 32], gene expression [30], and polyadenylation [33, 34]. Once trained, such models prioritize molecular quantitative trait loci (QTL) among non-functional variants and accurately estimate variant effect sizes [27, 30]. Previous work leveraged variant effect predictions (VEPs) from the Basenji model [23] to prioritize more than twice as many causal expression QTLs (eQTLs) in GTEx samples [35].

Since molecular phenotypes mediate trait heritability, we hypothesize that sequence-to-function models may provide insights into functional variant relevance in complex trait GWAS. While earlier work [36] found that sequence model annotations were not enriched for complex trait heritability conditioned on baseline annotations, it relied on older-generation models. Recent work demonstrated that ChromBPNet and Enformer VEPs were enriched for complex trait heritability, constructing a trait-agnostic variant-to-disease (V2D) score for prioritizing causal GWAS variants [37]. However, this approach does not perform functionally informed fine mapping—VEPs were applied post hoc by scaling PIPs after fine mapping rather than incorporated into SuSiE’s probabilistic model, thus not exploiting the full potential of jointly modeling GWAS data with functional priors. Rare variant association testing frameworks including DeepRVAT and Gruyere have further demonstrated the utility of Enformer VEPs in gene-based association tests [38]. Compared to Enformer, Borzoi estimates variant effects on a more diverse set of molecular phenotypes including gene expression, splicing, and polyadenylation by modeling RNA-seq in addition to ChIP-, CAGE-, DNase-, and ATAC-seq [30].

Here we present Sniff, a principled framework integrating Borzoi VEPs with 187 baseline-LF annotations using PolyFun to perform functionally informed fine mapping across 15 complex traits from the UK Biobank. Compared to fine mapping based solely on baseline-LF annotations (PolyFun-Baseline), Borzoi-informed fine mapping identifies 255 additional likely causal variants (9.45% increase) at PIP > 0.8. Sniff-prioritized variants demonstrate increased activity in Massively Parallel Reporter Assays (MPRAs) and exhibit >2-fold enrichment for expression-modifying variants relative to controls. For most traits, genes nominated by Sniff fine mapping rank higher by the similarity-based gene prioritization approach PoPS compared to genes nominated by PolyFun-Baseline. Sniff successfully nominates trait-relevant genes, such as *IL4R* for eczema and *PKN2* for diastolic blood pressure. Importantly, Sniff enables investigation of sequence determinants underlying fine-mapped variants through Borzoi, generating hypotheses about why particular variants are prioritized and how they mediate GWAS associations.

### 3 Results

### 3.1 Overview of Sniff fine mapping

We retrieved GWAS summary statistics for 15 complex traits in the UK Biobank (UKB), previously defined as maximally uncorrelated traits among 49 UK Biobank traits (Methods) [8]. We computed Borzoi variant effect predictions (VEPs) for 19,534,182 genotyped and imputed autosomal variants, including small indels (Methods). Borzoi predicts coverage at 32 bp resolution across 7,611 human functional genomics tracks including ChIP, ATAC/DNase, CAGE, and RNA-seq. To compute Borzoi VEPs, we compared predicted coverage for reference sequences to predicted coverage when the reference allele is replaced with the alternative. We used a bin-level L2 norm (“L2 score”) to compare predicted coverage vectors for each output track, resulting in a VEP vector with dimensionality 7,611 (Fig. 1A, Methods) [30]. Previous experiments demonstrated that large multitask models can suffer when predicting variant direction of effect on gene expression [39, 40]; our use of the unsigned L2 score focuses on variant effect magnitudes for prioritizing causal variants.

**Figure 1:**
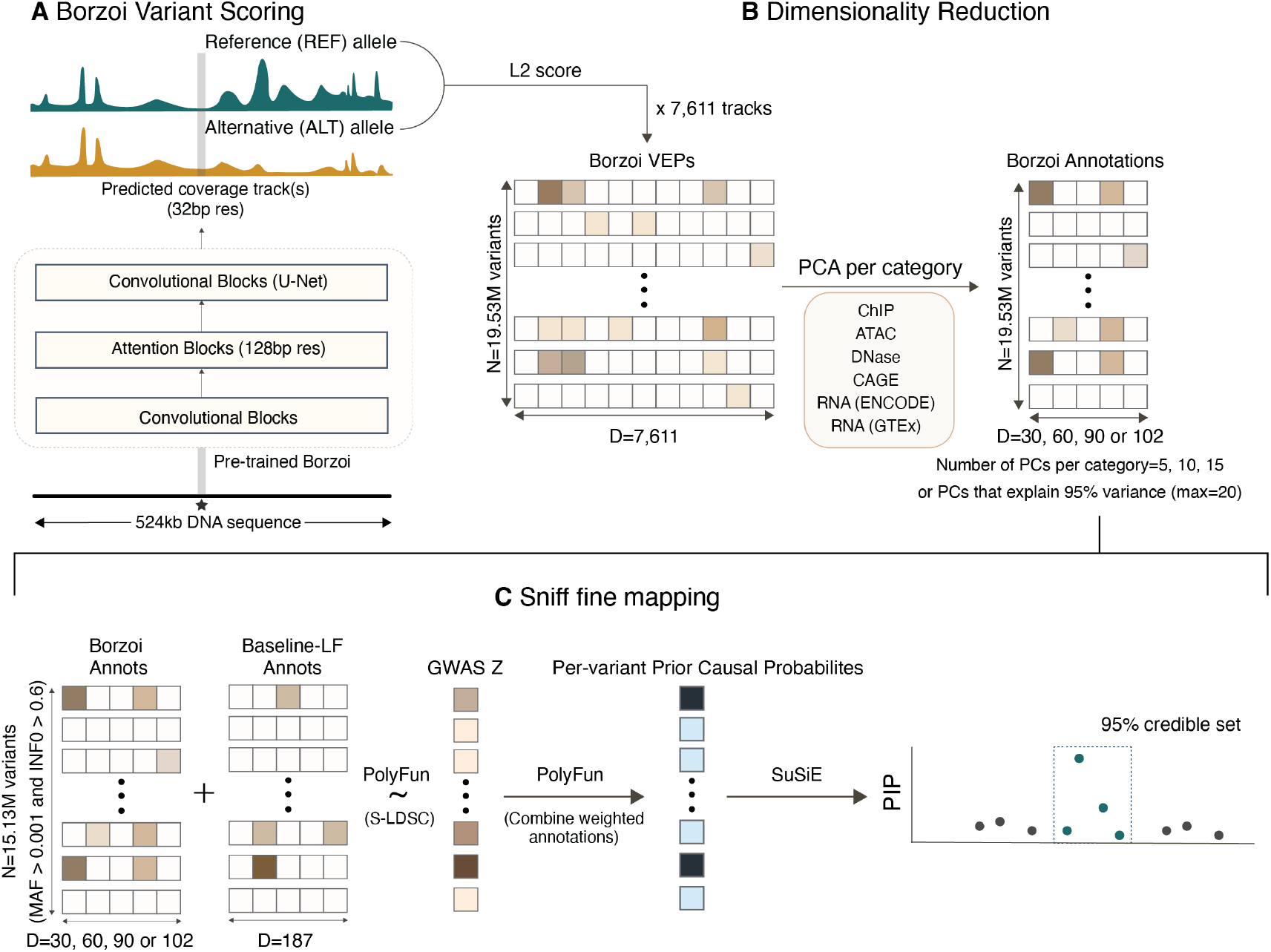
Sniff fine mapping overview. **A** The pre-trained Borzoi model estimates variant effects across 7,611 functional genomics coverage tracks using the L2 score. We computed Borzoi VEPs for ∼19.53 million common and low-frequency variants, resulting in a 19.53 million × 7,611 matrix. **B** A lower-dimensional representation of Borzoi VEPs is derived via PCA. We performed separate PCAs for each of six assay categories and retained either 5, 10, 15 PCs per category, or the number of PCs explaining 95% of variance (maximum 20 PCs per category). This yields 30, 60, 90, or 102 Borzoi VEP-derived PC annotations. **C** Borzoi annotations are combined with 187 baseline-LF(v2.2) annotations via PolyFun. PolyFun estimates prior causal probabilities for each SNP, which are then used within SuSiE for genome-wide fine mapping.

Computing indel VEPs with convolutional neural networks is challenging because indels introduce shifts in alternative sequences relative to reference sequences [41]. Sequence shifts cause misalignment in operations like max pooling and changes in output bin boundaries leading to artifactually inflated bin-level scores for indels compared to substitution single nucleotide polymorphisms (SNPs) (Supp. Fig. 1A) [41]. Because fine mapping jointly considers both indels and SNPs at genomic loci, biases in indel VEPs can introduce fine mapping biases. To mitigate inflation in indel VEPs, we computed indel VEPs using the “stitch” strategy that reduces differences between Borzoi SNP and indel VEP distributions (Methods, Supp. Fig. 1B) [41]. To fully ensure calibration, we quantile normalized the indel and SNP VEP distributions (Methods).

To integrate Borzoi VEPs into a Bayesian fine-mapping framework, we used PolyFun, which employs regularized stratified LD score regression (S-LDSC) [10, 42] to weight variant annotations given GWAS summary statistics and corresponding LD information [8]. Since high-dimensional correlated annotations would introduce regression instability, increase computational burden, and reduce interpretability, we performed principal component analysis (PCA) to reduce Borzoi VEP dimensionality before PolyFun integration (Methods). We divided Borzoi VEPs into six categories: ATAC, DNase, CAGE, ChIP, RNA (GTEx) and RNA (ENCODE), and performed PCA for each category (Methods). Since we did not know a priori the optimal number of PCs, we tested four configurations—retaining either 5, 10, or 15 PCs per category, or retaining PCs explaining 95% of variance (maximum 20 PCs per category). This resulted in reduced sets of 30, 60, 90, or 102 Borzoi-derived PCs per variant (referred to as Borzoi-30 through Borzoi-102 annotation sets, Fig. 1B).

We used PolyFun to derive variant prior causal probabilities given either only the 187 baseline-LF annotations (PolyFun-Baseline) or by combining BaselineLF annotations with Borzoi annotations (Sniff) (Fig. 1C). Prior to running PolyFun, we filtered variants to retain SNPs with MAF > 0.001 and INFO score > 0.6, resulting in 15,139,199 SNPs (Methods). Following the PolyFun strategy [8], we performed genome-wide fine mapping on 2,762 overlapping 3 Mb genomic windows using a 1 Mb stride (Methods). We used SuSiE to perform variant fine mapping [5, 6]. SuSiE outputs a PIP for each variant representing posterior causal probability. Because our fine mapping windows are tiled (variants may be fine mapped in up to three different windows), we assign variant PIPs based on the window where each variant is most central [8].

### 3.2 Evaluating Sniff fine mapping

To evaluate Borzoi VEPs’ contribution to variant prioritization, we compared Sniff fine mapping to PolyFun-Baseline and SuSiE-uniform (fine mapping without functional priors). Aggregated across 15 UKB traits, Sniff fine mapping increased the number of fine-mapped variants compared to the BaselineLF-only approach (Fig. 2A, Supp. Fig. 2A). At PIP> 0.5, incorporating the Borzoi-60 and Borzoi-102 annotations led to the largest gains relative to PolyFun-Baseline, with 394 (+7.44%) and 380 (+7.17%) additional fine-mapped variants, respectively (Supp. Fig. 2A). At PIP> 0.8, incorporating the Borzoi-102 annotation set yielded the greatest increase relative to PolyFun-Baseline, resulting in 255 (+9.45%) additional fine-mapped variants (Fig. 2A). At the same PIP threshold, Sniff (with Borzoi-102 annotations) resulted in 920 (+45.23%) additional fine-mapped variants compared to SuSiE-uniform. Based on these metrics, we selected the Borzoi-102 annotation set for all subsequent Sniff fine mapping.

**Figure 2:**
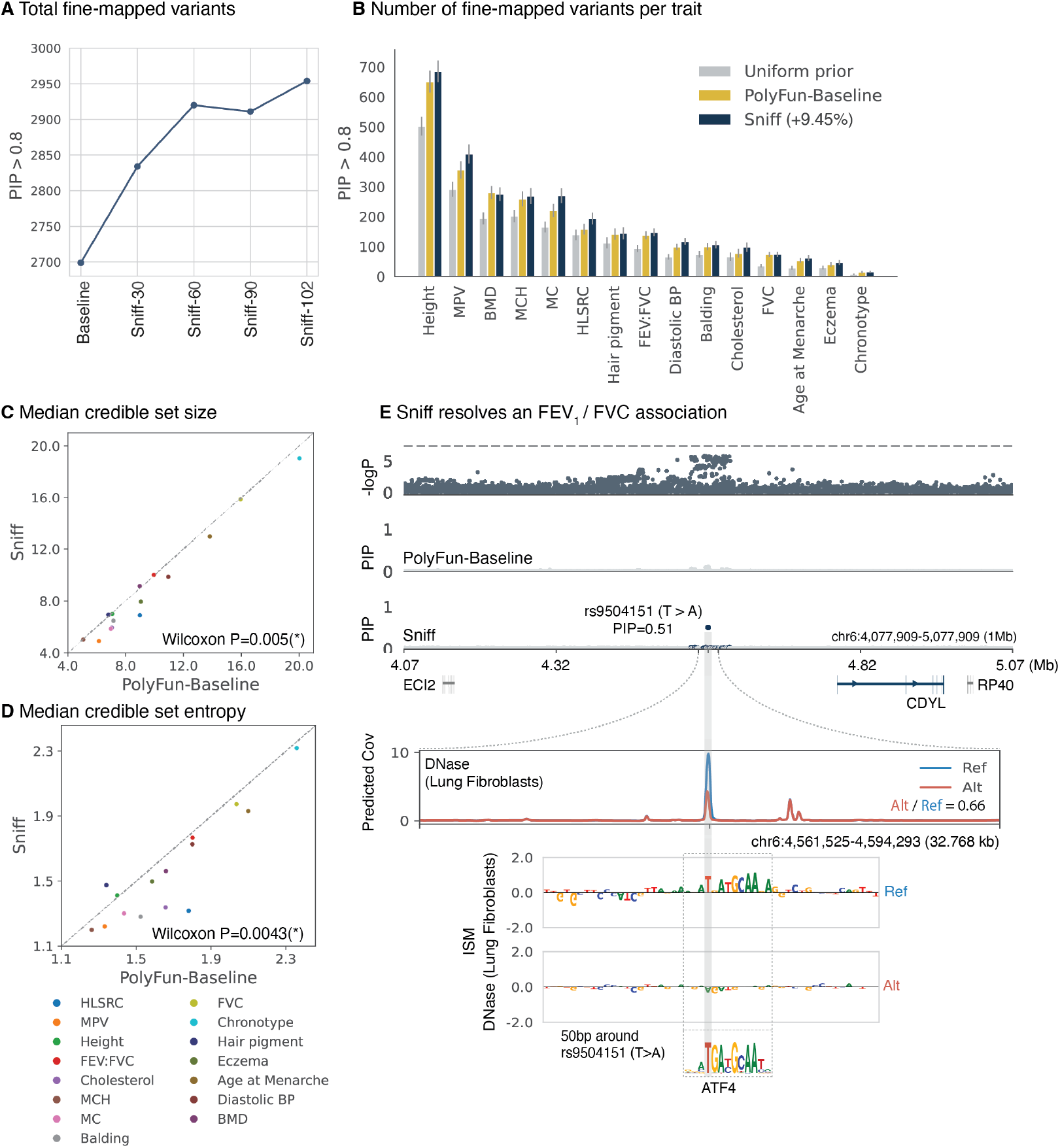
Evaluating Sniff fine mapping. **A** Integrating the Borzoi-102 annotation set with baseline-LF annotations (Sniff fine mapping) produces the largest gain in fine-mapped variants compared to baseline-LF-only approach at PIP>0.8. Counts are aggregated across 15 UKB traits. Sniff-30 through Sniff-102 refer to Sniff fine mapping with 30, 60, 90 and 102-dimensional Borzoi annotation sets respectively. All subsequent analyses use the Borzoi-102 annotation set. **B** Sniff fine mapping increases fine-mapped variants compared to BaselineLF-only approach for 13 of 15 traits (PIP > 0.8), yielding 255 additional fine mapped variants (9.45% increase). **C** Sniff fine mapping produces smaller median credible set sizes across 15 UKB traits (P=0.005, Wilcoxon signed-rank test). **D** Sniff fine mapping produces smaller median credible set entropy across 15 UKB traits (P=0.0043, Wilcoxon signed-rank test). **E** Example of an FEV_1_/FVC GWAS association resolved by Sniff. rs9504151 is fine mapped with PIP=0.51, implicating the *CDYL* gene. The polymorphism disrupts a putative ATF4 motif, resulting in decreased chromatin accessibility (DNase) at the locus in human lung fibroblasts.

When analyzed by trait, Sniff fine mapping increased fine-mapped variants compared to PolyFun-Baseline for 13 of 15 traits (PIP> 0.8) (Fig. 2B). Traits with the largest increases included cholesterol (+28.2%), reticulocyte count (+24.5%), monocyte count (+23%), and diastolic blood pressure (+17.3%). Traits showing the smallest increases included hair pigmentation (+2.8%) and lung forced vital capacity (FVC, no change). The only trait with decreased fine-mapped variants was bone mineral density (−2.5%), likely reflecting the limited bone-related Borzoi training datasets.

Beyond PIPs, SuSiE outputs 95% credible sets containing the likely causal variant with 95% probability. Credible set size evaluates fine mapping success; smaller sets indicate more precise causal variant localization. Across 15 UKB traits, Sniff fine mapping resulted in significantly smaller median credible set sizes (Wilcoxon signed-rank P=0.005, Fig. 2C). However, smaller credible set sizes do not necessarily imply greater causal variant certainty. For instance, a credible set with three variants each assigned PIP=0.33 is less informative than a credible set with four variants where one has PIP=0.8. To assess fine mapping certainty, we computed entropy of the variant PIP distribution within each credible set (Methods). Sniff fine mapping resulted in significantly decreased median credible set entropy across 15 UKB traits (Fig. 2D, Wilcoxon signed rank P=0.0043).

To illustrate Sniff’s ability to resolve previously ambiguous associations, we examined the *CDYL* locus. This locus associates with the FEV_1_ (forced expiratory volume) / FVC (forced vital capacity) ratio GWAS, although it falls marginally below the typical GWAS Bonferroni-derived significance threshold (5 × 10^*-*8^). In the genome-wide fine mapping, associations below global significance can potentially emerge from the Bayesian analysis when they have strong functional support. While neither SuSiE nor PolyFun-Baseline resolve this association, Sniff fine maps a single variant rs9504151 with PIP=0.51 near *CDYL* (Fig. 2E). Mouse experiments have shown disruption of *CDYL* impairs mouse lung epithelium differentiation and maturation [43], making it a plausible causal gene. Analyzing Borzoi attributions reveals rs9504151 disrupts a putative ATF4 binding motif, decreasing predicted chromatin accessibility and H3K27ac across multiple tissues. The greatest fold change decrease occurs in lung fibroblasts (Fig. 2E), suggesting this variant reduces enhancer activity, potentially regulating *CDYL* expression in lung fibroblasts. This example demonstrates how Sniff generates testable hypotheses explaining the functional basis of GWAS associations.

### 3.3 Functional properties of Sniff-prioritized variants

To investigate Sniff fine mapping, we extracted variants uniquely prioritized by Sniff. Sniff-prioritized variants were defined as variants assigned low PIPs by both SuSiE and PolyFun-Baseline (PIP *<* 0.1) that exhibited increased PIPs with Sniff (Methods). At ΔPIP (Sniff - PolyFun-Baseline) > 0.25, Sniff prioritized 1,158 unique SNPs. We first tested the Sniff-prioritized variant set for baseline annotation enrichment (Methods). Sniff-prioritized variants were significantly enriched for functional annotations including promoter-flanking regions (odds ratio=104.56), promoters (odds ratio=40.34), enhancers (odds ratio=30), and non-synonymous variants (odds ratio=26.8) (Methods, Supp. Fig. 3A). We next asked whether Sniff-prioritized variants were previously fine mapped in other traits. We found that 13.13% of Sniff-prioritized variants were previously fine mapped (PIP>0.1) in at least one other trait from 97 UKB-derived complex traits, compared to 2.69% of control variants from the same credible sets (Methods). At a more conservative ΔPIP (Sniff - PolyFun-Baseline) > 0.5, Sniff prioritized 299 unique SNPs, of which 20.07% were pleiotropic, compared to 4.7% of credible set-matched control SNPs (Methods, Supp. Fig. 3B). This is consistent with evidence that GWAS-discovered loci and underlying causal variants are highly pleiotropic [8, 44, 45].

To determine whether independent evidence supported variants uniquely prioritized by Sniff fine mapping, we turned to massively parallel reporter assays (MPRAs). MPRAs provide functional evidence of variant regulatory activity in vitro across cell types. Siraj et al. [46] profiled allele-specific activities of >200,000 variants using MPRAs across 5 cell-types, measuring activity differences (Δ log_2_ fold change) between alleles to identify expression-modifying variants (emVars). We leveraged this dataset to assess allele-specific activity of variants uniquely prioritized by Sniff (Methods). Variants prioritized by Sniff were more likely to exhibit allele-specific regulatory activity compared to matched control variants from the same credible sets (Wilcoxon P*<*0.001, Fig. 3A, Fig. 3C, Methods). At ΔPIP> 0.25, 30.9% of Sniff-prioritized variants measured in the MPRA were classified as emVars, compared to 9.7% of credible set-matched controls, representing a >2.5-fold enrichment (Fig. 3B). At ΔPIP> 0.5, 32.9% of Sniff-prioritized variants were emVars, compared to 13% of controls, representing a >2-fold enrichment (Fig. 3D). We next analyzed variants showing differential prioritization by Sniff—specifically, variants that may have been fine mapped by both PolyFun-Baseline and Sniff but were promoted or demoted by Sniff relative to PolyFun-Baseline (Methods). Sniff-promoted variants (Supp. Table 3) exhibited increased allele-specific activity compared to Sniff-demoted variants (Wilcoxon P*<*0.001, Fig. 3E). Furthermore, 28.9% of Sniff-promoted variants were emVars compared to 20.7% of demoted variants (Fig. 3F). As an example, we consider a credible set near the ZBTB7A gene, which contains a significant association for corpuscular hemoglobin. Sniff down-weights rs7254272 compared to PolyFun-Baseline and up-weights rs188955288. Of the 11 of 13 credible set variants tested in the MPRA study (including all variants with PIP> 0.01), rs188955288 had the highest allele-specific activity (Fig. 3G), illustrating a scenario where Sniff fine maps a variant consistent with its MPRA-estimated effect on gene expression.

**Figure 3:**
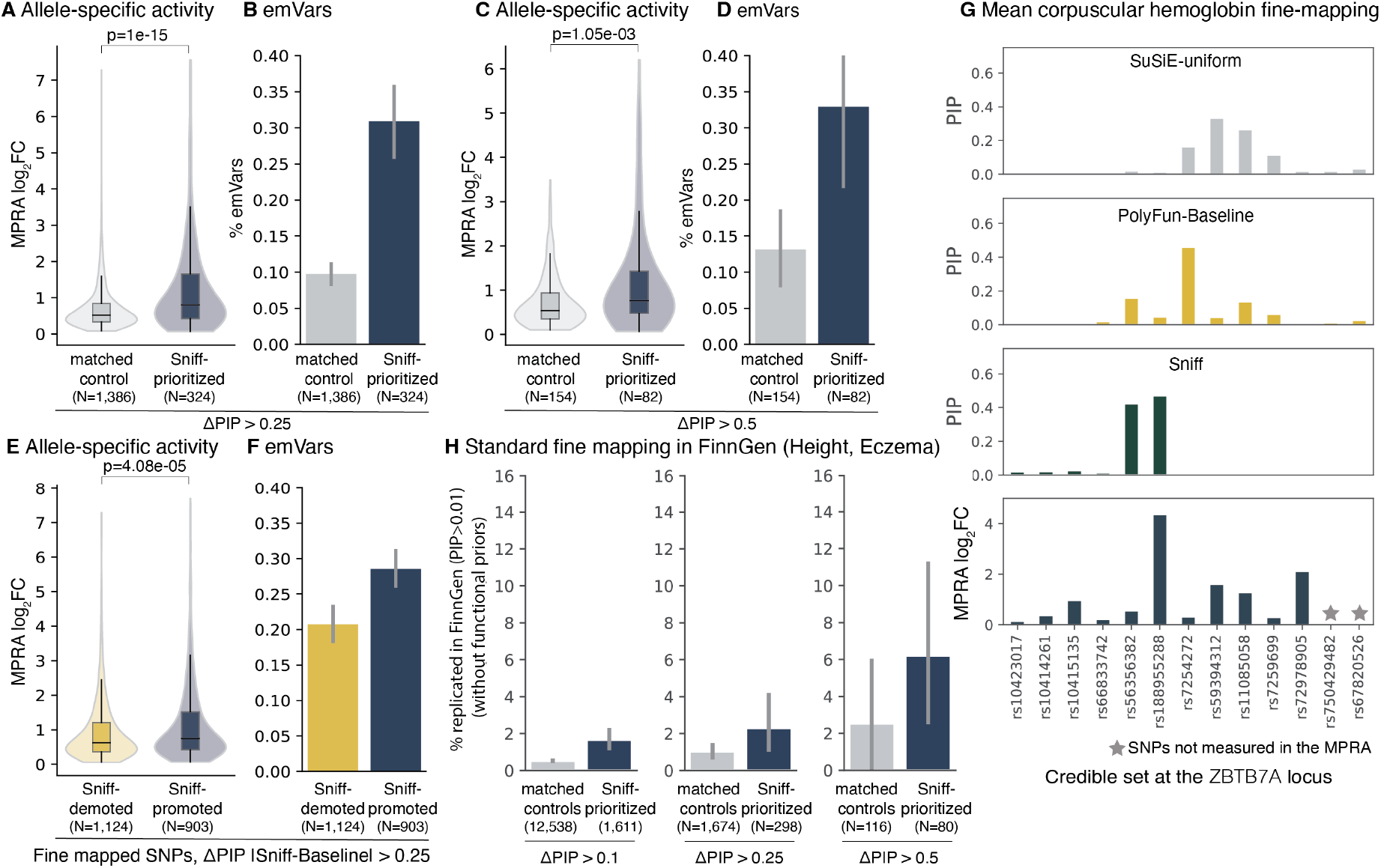
Investigating Sniff-prioritized variants. Measured allele-specific activity (MPRA log_2_ fold changes) averaged across five cell types and proportion of expression-modifying variants (emVars) for **A, B** Sniff-prioritized and credible set-matched controls (ΔPIP>0.25), **C, D** Sniff-prioritized and credible set-matched controls (ΔPIP>0.5), and **E, F** Sniff-promoted and Sniff-demoted variants (ΔPIP>0.25). **G** Distribution of credible set PIPs assigned by standard SuSiE, PolyFun-Baseline, and Sniff fine mapping at *ZBTB7A* locus in a mean corpuscular hemoglobin GWAS. Sniff uniquely promotes rs188955288 and rs56356382, of which rs188955288 has the highest measured allele-specific activity within this credible set. **H** Percentage of Sniff-prioritized and credible set-matched control variants prioritized in FinnGen with standard SuSiE without functionally informed priors.

To further validate the biological plausibility of Sniff-prioritized variants, we examined fine mapping results from the independent FinnGen cohort [47]. Due to LD pattern variation among populations, we hypothesized that variants prioritized by Sniff may also be recovered by standard fine mapping in FinnGen, even without functional priors. Using FinnGen fine mapping data for height and eczema GWAS, we observed that Sniff-prioritized variants (Sniff - Baseline PIP > 0.5) replicated at higher rates (6.25% with FinnGen PIP > 0.1) than control variants from the same credible sets (2.58%), despite FinnGen fine mapping being agnostic to functional annotations (Fig. 3H). When considering only variants with MAFs>0.001 in the FinnGen cohort, 8.77% of Sniff-prioritized variants were fine mapped in FinnGen (PIP>0.1) compared to 2.7% of credible-set matched control variants (Supp. Fig. 3C), supporting Sniff’s ability to prioritize variants with greater likelihood of being truly causal.

### 3.4 Borzoi annotations are enriched for trait heritability

To investigate the basis of Sniff fine mapping, we asked whether Borzoi annotations were enriched for trait heritability—specifically, whether variants with high VEPs in specific tissues or cell types were enriched for trait heritability. Since we use Borzoi-derived PCs as annotations, we first sought to interpret the PCs. We focused on PCs derived from Borzoi VEPs on RNA-seq coverage across 31 GTEx tissues, represented by 87 functional tracks (Methods); 13 PCs collectively explain 95% of VEP variance across these 87 tracks. Examining principal component loadings revealed that PCs capture both shared and tissue-specific effects and recapitulate biologically meaningful similarities between tissue types (Fig. 4A). PC1 shows uniformly distributed loadings across the 87 GTEx RNA-seq VEP vectors, suggesting it captures shared regulatory activity (Fig. 4A). PC2 captures variation in liver, pancreas, small intestine, and stomach VEPs; PC3 captures variation in muscle, heart, and liver VEPs; and PC4 captures variation in spleen and blood VEPs, representing effects on blood and immune-specific gene expression (Fig. 4A). Several PCs exhibit both positive and negative loadings. For example, PC9 assigns positive weights to heart VEPs and negative weights to muscle VEPs, i.e. variants with high PC9 scores tend to have stronger predicted effects in heart, while those with low scores show stronger effects in muscle (Fig. 4A). This pattern may reflect shared regulatory architecture between heart and muscle, with PC9 capturing a regulatory axis distinguishing cardiac- and muscle-specific variant effects.

**Figure 4:**
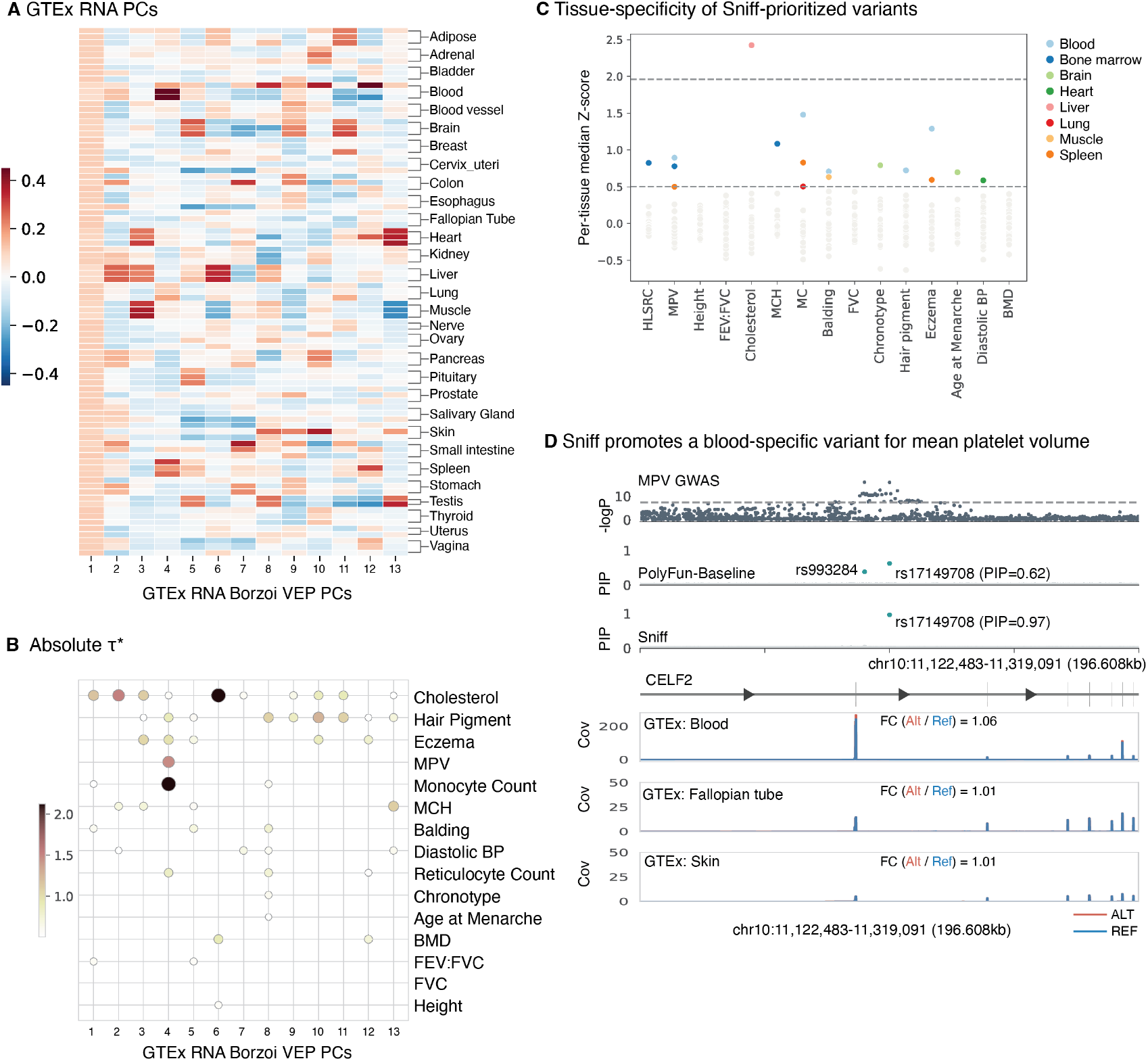
Borzoi-derived annotations are enriched for trait heritability. **A** Loadings for 13 PCs explaining 95% of variance across 87 GTEx RNA-seq VEPs. **B** Absolute values of standardized effect sizes *τ* ^*∗*^ for each Borzoi PC, computed by S-LDSC conditioned on 187 BaselineLF annotations. |*τ* ^*∗*^| values > 0.5 are plotted. Dot size and color correlate with |*τ* ^*∗*^|value. **C** Z-scores of Sniff-differential variant L2 scores are computed in each tissue relative to L2 scores across 31 GTEx tissues included in the Borzoi training data. Median Z-scores are plotted for each of 31 tissues for each trait. Tissues where median *Z* > 0.5 are labeled; tissues where median *Z <* 0.5 are colored grey. Dashed lines correspond to Z=0.5 and Z=1.96. **D** Example variant rs17149708 is predicted to have blood-specific effects and is promoted by Sniff fine mapping in the mean platelet volume GWAS. The three tissues (among 31 GTEx tissues) with the largest predicted fold change (Alt/Ref) are plotted, with the largest predicted effect on *CELF2* expression observed in GTEx blood. Fold change is computed over a 196,608 base pair window surrounding the variant. The blue track represents reference allele predictions; the red track represents alternative allele predictions.

We applied S-LDSC to estimate whether Borzoi VEP-derived PCs were enriched for trait heritability [10, 42]. We assessed whether PCs contributed to trait heritability after conditioning on the 187 baseline-LF annotations (Methods). For each of the 15 UKB traits, we computed the standardized effect size *τ* ^*∗*^ for each PC. Following prior work [36, 37], we considered annotations with absolute *τ* ^*∗*^ > 0.5 to be enriched for heritability and likely to contribute to disease risk. SinceB Borzoi PCs can take both positive and negative values, we observe both positive and negative *τ* ^*∗*^ estimates (Supp. Fig. 4). Because each PC assigns positive and negative weights to tissue VEPs, we use the absolute *τ* ^*∗*^ estimate to assess PC relevance and interpret signed *τ* ^*∗*^ estimates in context of PC loadings. We find that Borzoi-derived annotations are highly enriched for trait heritability (Fig. 4B). PC6, associated with liver VEPs, is highly enriched for cholesterol heritability (*τ* ^*∗*^ = 2.1), followed by PC2 (associated with liver, pancreas, intestine and stomach RNA VEPs, *τ* ^*∗*^ = 1.5) and PC3 (associated with liver, muscle and heart VEPs, *τ* ^*∗*^ = 1.1). PC4 (associated with blood and spleen VEPs) is enriched for heritability across blood and immune-related traits including monocyte count (*τ* ^*∗*^ = 2.1), mean platelet volume (*τ* ^*∗*^ = 1.5), eczema (*τ* ^*∗*^ = 0.98) and high light scatter reticulocyte count (*τ* ^*∗*^ = 0.84), whereas PC13 (associated with heart VEPs) is enriched for diastolic BP heritability (*τ* ^*∗*^ = 0.62).

Next, we directly analyzed the predicted tissue specificity of variants differentially prioritized by Sniff (Methods). To quantify tissue specificity, we calculated Z-scores for variant-tissue pairs relative to each variant’s predicted effect across 31 GTEx tissues (Methods). We observed that Sniff-prioritized variants are predicted to have tissue-specific activity in trait-relevant tissues: cholesterol variants were most likely to be liver-specific (Z=2.42), consistent with the liver’s role in lipid metabolism (Fig. 4C). Hemoglobin and reticulocyte count variants were most likely to be bone marrow-specific (Z=1.08, Z=0.78, respectively), monocyte count variants were most likely to be blood (Z=1.48) and spleen-specific (Z=0.50), and diastolic blood pressure variants were most likely to be heart specific (Z=0.59, Fig. 4C). However, consistent with S-LDSC-derived enrichment, for a subset of traits, Sniff-prioritized variants did not point to a single trait-relevant tissue. Several explanations may account for this observation: (1) absence of disease-relevant tissues in the Borzoi data (e.g., bone tissue for bone mineral density); (2) inherently multi-tissue phenotypes (such as height); or (3) limitations in the model’s capacity to detect tissue-specific effects. These observations suggest that integrating additional tissues and cell types into the model will enhance its ability to identify trait-relevant variants.

Figure 4D presents an example of a variant predicted to have tissue-specific activity and promoted by Sniff at the *CELF2* locus, which is significantly associated with mean platelet volume. PolyFun-Baseline identifies a credible set with two variants (rs17149708, PIP=0.62 and rs993284, PIP=0.38), whereas Sniff resolves the association to a single variant rs17149708 with PIP=0.97. At this SNP, the alternative allele is predicted to increase *CELF2* expression, with the largest increase observed in GTEx blood samples among the 31 GTEx tissues, aligning with the expected relevance of blood gene expression for platelet volume GWAS.

### 3.5 Sniff identifies trait-relevant genes

For associations in noncoding regions, the most common approach for prioritizing causal genes is the distance-based “nearest gene” approach, where the gene nearest to a lead variant is nominated as a target gene [48]. In contrast to genes nominated based on lead variants, genes nominated based on distance to causal variants are more likely to influence the trait [48]. However, not all associations can be fine mapped to a single causal variant. Instead, genes nearest to 95% credible sets, weighted by variant PIPs, can be nominated as target genes [48](Methods). In this framework, gene discovery depends on the underlying fine mapping. This is illustrated in Fig. 5A, where differential credible set discovery can lead to differential gene assignment. To assess the utility of Sniff fine mapping, we nominated target genes based on their weighted distance to Sniff-derived credible sets for each of the 15 UKB traits (Methods). To benchmark Sniff-based gene discovery, we additionally nominated genes based on PolyFun-Baseline and SuSiE-uniform fine mapping.

**Figure 5:**
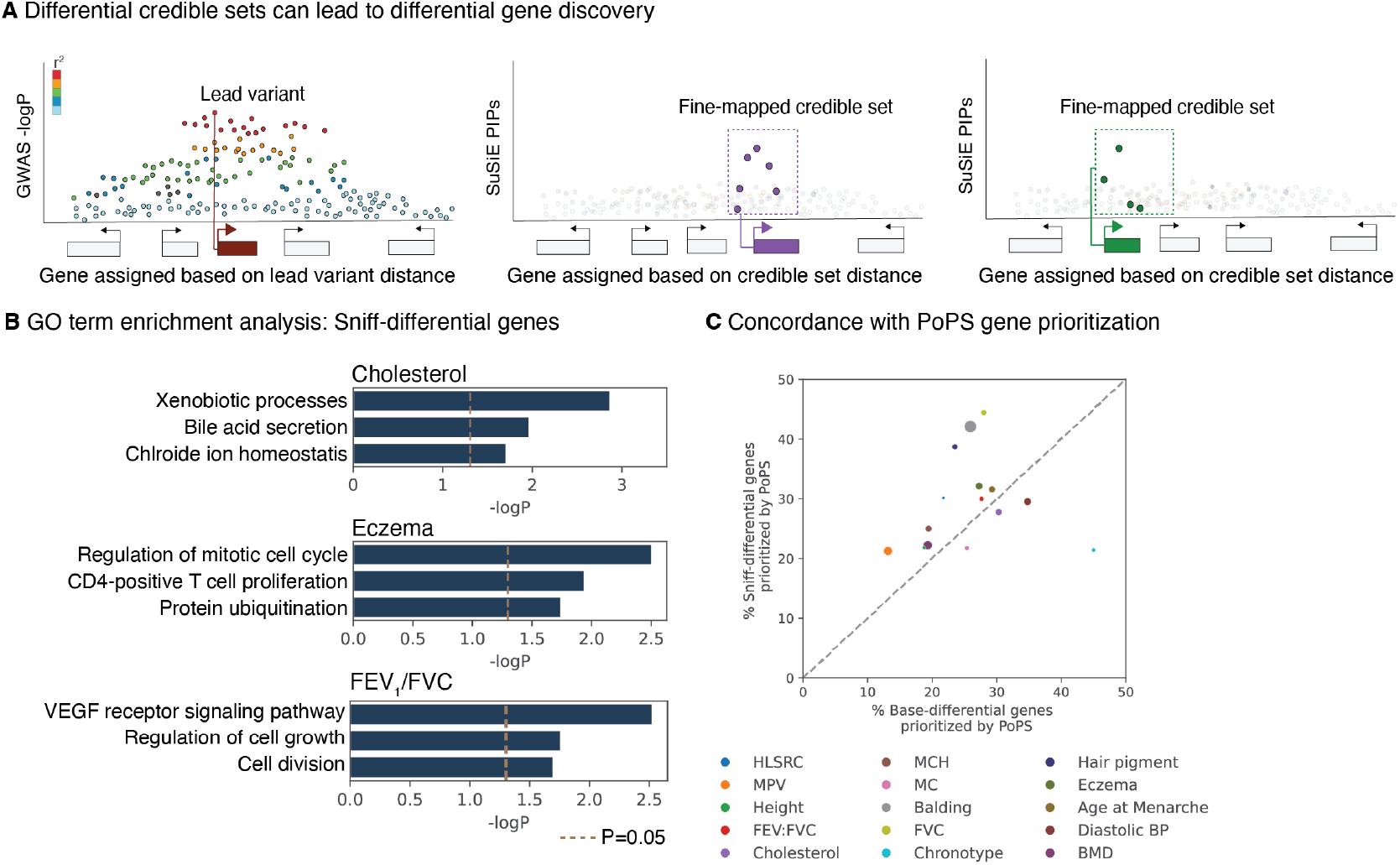
Sniff fine mapping augments gene prioritization. **A** Schematic illustrating how differentially fine-mapped credible sets can nominate distinct genes. **B** Gene Ontology (GO) term enrichment analysis of Sniff-differential genes for selected traits (cholesterol, eczema, reticulocyte count and balding). **C** Percentage of Sniff-differential genes prioritized by PoPS compared to baseline-differential genes prioritized by PoPS. Dot size is proportional to the number of credible sets discovered (equivalent to number of genes nominated) by standard fine mapping.

We compared genes nominated by Sniff fine mapping to genes nominated by PolyFun-Baseline fine mapping (Methods). The gene sets prioritized by the two approaches are similar, with a median Jaccard index of 0.81 across the 15 traits (Supp. Fig. 5A). 89.43% of gene-trait pairs nominated by Sniff fine mapping were also nominated by PolyFun-Baseline, and conversely, 89.37% of gene-trait pairs nominated by PolyFun-Baseline were also nominated by Sniff (Supp. Fig. 5B). Compared to PolyFun-Baseline, Sniff fine mapping differentially identified 869 gene-trait pairs across the 15 traits, of which 34.3% were also identified by SuSiE-uniform fine mapping. We focused on this set of Sniff-differential genes and asked whether Sniff fine mapping discovers trait-relevant genes. To assess trait relevance, we first performed Gene Ontology (GO) enrichment analysis—we find that genes differentially identified by Sniff are enriched for trait-relevant GO terms (Fig. 5B). For example, Sniff-differential genes prioritized for eczema are enriched for immune cell proliferation including “regulation of mitotic cell cycle” and “CD4+ T cell proliferation”, whereas Sniff-differential genes prioritized for cholesterol are enriched in terms related to cholesterol metabolism including “xenobiotic processes” and “bile acid secretion”.

To quantitatively assess Sniff-based gene prioritization, we asked how likely Sniff-differential genes were to be nominated by orthogonal methods compared to genes differentially prioritized by PolyFun-Baseline. We used the Polygenic Priority Score (PoPS), a gene prioritization method that considers trait-relevant gene features from thousands of features including single-cell RNA-seq, protein-protein interactions (PPI), and gene annotations [49]. The gene with the highest PoPS score in a 1 Mb window around a fine-mapped credible set is nominated as the target (Methods). We ran PoPS for the 15 UKB traits and analyzed PoPS scores for Sniff-differential genes. Sniff-differential genes were ranked higher by PoPS (average median rank = 2.9 across traits) compared to baseline-differential genes (average median rank = 3.23 across traits). On average across the 15 UKB traits, 29.3% of Sniff differential genes were the highest-ranked genes in 1 Mb loci around fine-mapped variants, compared to 26% of baseline-differential genes (Fig. 5C). The largest gains were seen for traits FVC (44.5% Sniff-differential genes ranked first, compared to 28% baseline-differential genes), high light scatter reticulocyte count (30.1% Sniff-differential genes ranked first, compared to 21.7% baseline-differential genes), and balding (42.1% Sniff-differential genes ranked first, compared to 25.9% baseline-differential genes). A notable exception was chronotype, where 21.4% of Sniff-differential genes were ranked 1 by PoPS compared to 45% of PolyFun-Baseline-differential genes. For chronotype, both PolyFun-Baseline and Sniff identified relatively few high PIP variants, potentially due to reduced power from a smaller GWAS sample size (N=155,100), suggesting that Sniff fine mapping may provide limited benefits for poorly-powered GWAS.

Examples of trait-relevant Sniff-differential genes include *SPRED2, GSTA1*, and *TP53INP1* for cholesterol; *IL4R, SCML4*, and *SESN1* for eczema; and *DUSP5, UBTD1*, and *LAPTM5* for monocyte count. The complete set of Sniff-differential genes is provided in Supplementary Table 4.

### 3.6 Sniff facilitates interpretation of molecular mechanisms driving trait associations

To demonstrate Sniff fine mapping’s utility, we examine three uniquely identified genes. First, we focus on a diastolic blood pressure association marginally below the GWAS significance threshold (rs7545929, GWAS p=5.14 × 10^*-*8^). Neither SuSiE nor PolyFun-Baseline identified a credible set at this locus; however, Sniff fine maps rs12041762 with PIP=0.64 (Fig. 6A), 508 kb from the nearest protein-coding gene *PKN2. PKN2* was previously implicated in systolic blood pressure and hypertension GWAS, experimental studies demonstrated its role in blood pressure regulation, and it is the highest scoring PoPS gene in a 2Mb window around the variant (Fig. 6D). Model attributions computed over the region centered on rs12041762 show that the alternative allele leads to creation of a GATA2 motif, leading to increased predicted H3K4me1 in heart tissue (Fig. 6A), suggesting that rs12041762 likely drives the observed GWAS association by regulating *PKN2* expression levels

**Figure 6:**
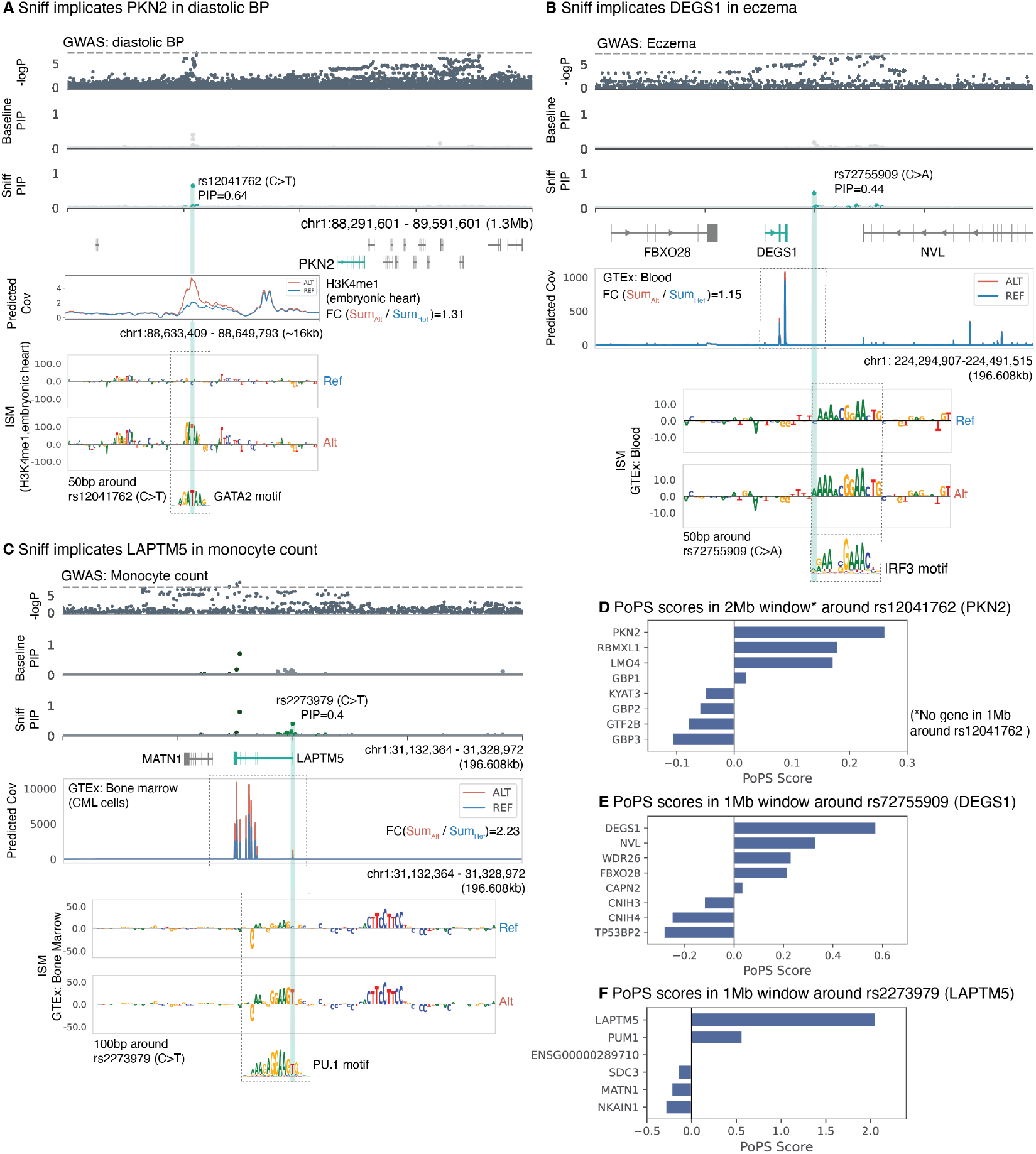
Borzoi-based fine mapping enables interpretation of molecular mechanisms driving trait associations. **A** Sniff uniquely fine maps rs12041762 with PIP=0.64 in a diastolic BP GWAS. In silico mutagenesis (ISM) reveals the ALT allele (T) leads to creation of a putative GATA2 motif, predicted to increase H3K4me1 in embyonic heart tissue. **B** Sniff uniquely fine maps rs72755909 with PIP=0.44 in an eczema GWAS. ISM performed in whole blood reveals the ALT allele (A) leads to a predicted increase in IRF3 motif strength and associated increase in *DEGS1* expression in GTEx whole blood. **C** Sniff discovers an additional credible set at the LAPTM5 locus, fine mapping rs2273979 with PIP=0.4 in a monocyte count GWAS. ISM performed a in CML cells reveals the ALT allele (T) leads to predicted increase in PU.1 motif strength and an associated predicted increase in *LAPTM5* expression. For **A, B** and **C**, strongest motif matches from the JASPAR database are plotted. **D** PoPS scores for genes in a 1 Mb window around rs2273979 and rs72755909. Because rs12041762 implicates PKN2, outside a 1 Mb window centered on the variant, PoPS scores are plotted for all genes in a 2 Mb window.

Next, we examine an eczema association at the *DEGS1* locus. This locus represents another sub-significant association. Sniff uniquely fine maps rs72755909 with PIP=0.44 (Fig. 6B). rs72755909 lies downstream of the DEGS1 gene and is predicted by Borzoi to modulate *DEGS1* expression (Fig. 6B). *DEGS1* is highly expressed in skin tissue and is associated with sphingolipid metabolism, pathway involved in maintaining skin barriers. The largest predicted effect on gene expression is in GTEx blood, with a smaller but significant effect in GTEx skin. Based on model attributions, the variant is in the flanking region of an IRF3 motif, and the alternative allele is expected to increase motif strength, consistent with the predicted increase in *DEGS1* expression (Fig. 6B). Although *DEGS1* is the highest-scoring PoPS gene at this locus (Fig. 6E), we note that PoPS would not have nominated *DEGS1* since no lead variant or credible set was discovered in this region by standard fine-mapping approaches. Sniff fine mapping therefore enables novel gene discovery via PoPs by augmenting causal variant fine mapping.

Finally, we examine *LAPTM5*, a gene implicated by Sniff fine mapping in the monocyte count GWAS. Both SuSiE and PolyFun-Baseline result in a single fine-mapped credible set, closest to the transcription start site of *MATN1. MATN1* is also implicated by the lead variant. Sniff fine mapping discovers an additional credible set, fine mapping rs2273979 with PIP=0.4 (Fig. 6C). rs2273979 is 47 base pairs upstream of the *LAPTM5* gene and lies within a PU.1 motif. The largest effect on *LAPTM5* expression is predicted to be in GTEx bone marrow-derived CML cells, where the alternative allele is predicted to increase *LAPTM5* expression (Fig. 6C). Model attributions show the alternative allele leads to predicted increase in PU.1 motif strength (Fig. 6C). *LAPTM5* is a lysosomal membrane protein highly expressed in several immune cell-types including monocytes, macrophages, and B cells [50]; it negatively regulates lymphocyte activation and positively regulates macrophage signaling [51]. Within the 1 Mb region surrounding rs2273979, *LAPTM5* is assigned the highest PoPS score, supporting its candidacy as an effector gene (Fig. 6F). Sniff fine mapping thus nominates *LAPTM5* as a likely mediator of the genetic association with monocyte count, consistent with its immune regulatory functions.

## 4 Discussion

In this work, we present Sniff, a functionally informed fine-mapping framework that integrates Borzoi-derived variant effect predictions (VEPs) with broad baseline-LF functional annotations to prioritize causal variants in complex trait GWAS. Across 15 UK Biobank traits, Sniff increases fine mapping resolution. Sniff-prioritized variants are more than 2-fold enriched for expression-modifying activity in MPRAs, exhibit tissue-specific effects relevant to their associated traits, and nominate genes that exhibit greater consistency with orthogonal prioritization methods such as PoPS compared to fine mapping with baseline-LF annotations alone.

In contrast to the V2D framework, which applies functional score scaling post-hoc to fine map ping results, Sniff incorporates Borzoi-derived VEPs as priors within a Bayesian fine-mapping framework. This integration provides natural regularization—variants increase in priority only when both functionally plausible and statistically supported by GWAS data, preventing spurious prioritization of variants lacking population-level evidence. In contrast to generic, trait-agnostic priors, we leverage PolyFun’s trait-specific annotation weighting, allowing Sniff to derive context-dependent priors that identify and promote tissue-relevant variants. For example, cholesterol-associated variants prioritized by Sniff are predicted to have liver-specific effects, whereas variants prioritized by Sniff in blood-related phenotypes are predicted to have blood-specific effects, demonstrating biological coherence.

Sniff successfully prioritizes gain-of-function variants that conventional annotation-based approaches miss. These variants typically lie outside reference genome annotations yet have functional consequences detectable by sequence-based models. Additionally, our examples highlight cases where Sniff prioritizes variants that subtly modify transcription factor motif strength rather than completely disrupting binding, with modest predicted effects on gene expression. This capability aligns with emerging evidence that GWAS predominantly identifies variants with modest effects on important genes, rather than dramatic loss-of-function variants [52, 53].

This work has several limitations that present future opportunities. Borzoi’s training corpus, while extensive, remains incomplete at cell type resolution. If a trait-relevant cell type is missing from the model’s training data, the model may not identify variants that have effects in only that cell type. As an illustration, the limited benefit from incorporating Borzoi annotations when fine mapping the bone mineral density GWAS likely reflects sparse bone-related datasets in Borzoi, preventing the model from learning bone-specific regulation. We expect fine-mapping power to increase further across traits as the community builds models trained on additional trait-relevant cell type-specific chromatin accessibility and gene expression data [27, 54]. While training large multitask models on new cell types is compute-intensive, lightweight transfer learning to relevant single-cell datasets enables models to learn more comprehensive regulatory lexicons [55]. Several methods have been developed that transfer Borzoi to single-cell atlases or train on combined bulk and single-cell data including Borzoi Prime [56], Scooby [54] and Decima [57]. Since a significant proportion of trait heritability is expected to lie in cell type specific regulatory regions [17, 58], we expect that leveraging these models to derive VEPs would boost fine-mapping power.

Second, although Borzoi accurately predicts diverse regulatory processes at transcriptional and post-transcriptional levels, incorporating VEPs from models trained on regulatory processes not explicitly modeled by Borzoi such as translation initiation [59] and RNA half-life [60], as well as from missense variant effect predictors [61, 62] may further strengthen causal variant fine mapping. Finally, PolyFun’s reliance on S-LDSC assumes linear, additive annotation effects without interactions. Leveraging nonlinear functions to estimate per-variant prior causal probabilities could enhance the use of rich, high-dimensional VEPs from large multitask models such as Borzoi [63]. The recent KGWAS approach demonstrated how deep neural networks can aggregate annotations over large knowledge graphs to enhance GWAS discovery power [64]. We anticipate that applying similar strategies to aggregate Borzoi-derived VEPs will further improve fine mapping resolution and accuracy.

Sniff represents the first principled integration of allele-specific ML predictions into biobank-scale Bayesian fine mapping, establishing a framework for incorporating diverse molecular predictions into genetic association studies. As sequence-to-function models continue improving in accuracy and scope, we anticipate they will become standard tools for post-GWAS analysis, enabling mechanistic interpretation of genetic associations across the allelic frequency spectrum.

## 5 Methods

### Borzoi variant effect predictions

Borzoi is a hybrid convolution and transformer model that takes as input a one-hot encoded DNA sequence of length 524 kb and predicts coverage for 6,385 datasets including RNA-seq, CAGE, ATAC-seq, DNase-seq, and ChIP-seq. For a subset of RNA-seq and CAGE datasets, Borzoi predicts coverage on both positive and negative strands, leading to coverage predictions for a total of 7,611 tracks. Borzoi’s architecture and training procedure are described in detail in Linder et al [30].

Briefly, Borzoi predicts coverage in 32 bp bins; i.e., for an input sequence length 524 kb, Borzoi predicts coverage vectors of length 524 kb / 32 bp = 16,375. Let **x** ∈ {0, 1}^524,000*×*4^ represent a one-hot encoded input sequence and ℳ represent the Borzoi model. Borzoi outpu s coverage across 7,611 tracks, resulting in output **y** = ℳ(**x**) where **y** ∈ [0, ∞)^16,375*×*7,611^.

To score a variant, we extract the 524 kb sequence centered on the reference allele and compute model predictions **y**^REF^. We create an alternative sequence by replacing the reference allele with the alternative allele and recompute model predictions **y**^ALT^. We have previously observed that Borzoi predictions become less accurate as we approach the input sequence boundaries. Prior to computing variant effects, we crop Borzoi predictions, focusing on coverage predictions over 196,608 bp or 6,144 bins centered on the variant.

For the subset of datasets for which Borzoi predicts coverage on both positive and negative strands, we concatenate the stranded coverage predictions when computing variant scores, allowing us to consider each strand independently and compute an aggregate score over both strands. Let *t* be an index into the combined track dimension, *t* ∈ {1, 2, …, 6385}. We define a per-track variant score *s*_*t*_ as the *𝓁*_2_ norm across the length axis between log-transformed reference versus alternative coverage vectors. For each track, the variant score **s**_*t*_ is:

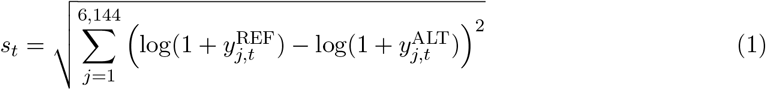

We compute a score independently for each output track *t*, such that **s** ∈ [0, ∞)^6,385^. The Borzoi training procedure applies a square root and scaling transformation to the coverage tracks **y** during training. Prior to computing the *𝓁*_2_ norm, we therefore perform an inverse transformation on the predicted **y**^REF^ and **y**^ALT^, allowing us to operate in the read count space.

Variant effect scores were computed using https://github.com/calico/baskerville/blob/main/src/baskerville/scripts/hound_snp.py. The *𝓁*_2_ score between log-transformed coverage vectors as described here can be computed using the ‘logD2’ option. We used a single Borzoi replicate to score all variants, accessed via https://storage.googleapis.com/seqnn-share/borzoi/f0/model0_best.h5. The parameter file required to score variants was accessed via: https://storage.googleapis.com/seqnn-share/borzoi/params.json. Variants were scored with the settings: *“--float16 -u --stats logD2 --shifts 0 --rc --indel stitch --require gpu”*.

### Borzoi variant effect prediction for indels

The 524 kb reference and alternative Borzoi input sequences are exactly aligned when scoring a substitution SNP, except at the variant position. Model predictions for the two sequences can therefore be compared at bin-level in an unbiased fashion. In contrast, the introduction of insertions and deletions (indels) leads to the alternative sequence becoming offset from the reference sequence. The alternative sequence is then aligned with the reference sequence either only on the left (before the indel) or only on the right (after the indel). Due to boundary-dependent operations like max pooling and the output bin comparison, misalignment between reference and alternative sequences leads to biased L2 scores for indels [41]. To overcome biases in indel variant effect scoring, we recently proposed the *shift augmentation stitching* strategy, which alleviates the indel bias [41]. We scored 1,907,577 indels using the stitching strategy. Observing that the distribution of L2 scores for SNPs still differed from the distribution of L2 scores for indels, we additionally performed quantile normalization to align their distributions.

### Sequence attributions with in silico mutagenesis

In silico mutagenesis (ISM) is used to attribute Borzoi’s prediction to individual nucleotides. Given an input genomic sequence *g*, ISM measures the sensitivity of model predictions to single nucleotides *i* within this sequence *g*. Nucleotide importance ISM_*i*_ is defined as the average change in model predictions when a nucleotide at position *i* is mutated to one of three potential nucleotides. Based on model definitions provided in the above section, let **y** = ℳ (**x**) where **y** ∈ [0, ∞)^16,375*×*7,611^. Let *g*_*mut,i,k*_ represent the sequence when the nucleotide at position *i* is mutated to one of three potential nucleotides, indexed by *k*. Then, nucleotide importance ISM_*i*_ for an output track *t* is defined as:

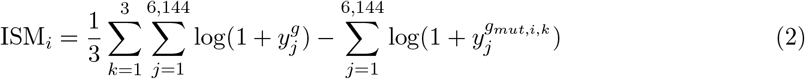

We used the implementation provided in the Baskerville GitHub repository to compute ISMs for sequences containing the reference and alternative alleles https://github.com/calico/baskerville/blob/main/src/baskerville/scripts/hound_ism_snp.py. Sequences were centered on the variant and ISM scores were computed for 100 nucleotides surrounding the variant of interest.

### Iterative principal component analysis (iPCA)

Borzoi estimates variant effects across 6,385 datasets. We categorize these datasets into six categories representing CAGE, DNase-seq, ATAC-seq, ChIP-seq, RNA-seq (ENCODE), and RNA-seq (GTEx), containing 638, 674, 232, 3,886, 868, and 87 VEPs per variant respectively. For each category, we perform an iterative principal component analysis (iPCA) on the resulting matrices of 19,534,182 variants by *p* VEPs, with *p* representing the number of datasets in that category. Rather than performing PCA on the entire dataset, incremental PCA processes data in batches and performs a partial SVD on each batch, allowing for an efficient approximation of principal components without needing to compute the full covariance matrix [65, 66]. In contrast to PCA, iPCA can efficiently scale to the large 19.53 million×*p* VEP matrices. We use the scikit-learn implementation of an iterative PCA algorithm and set the batch size to 100,000 variants [66]. We retain 5, 10, 15, or a number of principal components that explain up to 95% of the variance (for a maximum of 20 PCs) for each category.

### Summary statistics for UK Biobank-derived complex traits

We downloaded GWAS summary statistics for 49 UK Biobank-derived complex traits, made available by Weissbrod et al. (2020), https://alkesgroup.broadinstitute.org/polyfun_results. These summary statistics have since been moved to https://console.cloud.google.com/storage/browser/broad-alkesgroup-public-requester-pays. Summary statistics were derived from British-ancestry individuals within the UK Biobank, with an average sample size of 318,000 across the 49 traits. The sample sizes for each trait are available via PolyFun.

The summary statistics were computed using BOLT-LMM v.2.3.3 as described in Weissbrod et al. (2020), over 19,534,182 genotyped and imputed variants. From the set of 49 traits, we selected a subset of 16 maximally uncorrelated traits to analyze [8]. We excluded the trait “number of children” as it contains no fine mapped SNPs (PIP> 0.95) for either SuSiE or PolyFun run with the BaselineLF annotation set (PolyFun-Baseline) [8], resulting in a reduced set of 15 traits.

### Functionally informed fine mapping

We filtered the set of genome-wide genotyped and imputed autosomal SNPs to 15,300,176 SNPs with *MAF* ≥ 0.001 and INFO score ≥ 0.6. We additionally excluded the MHC region (chr6:25.5Mb– 33.5Mb), as well as two additional long-range LD regions (chr8:8Mb–12Mb, chr11:46Mb–57Mb) for all steps in the functionally informed fine mapping pipeline including computing LD scores and running PolyFun, S-LDSC, and fine mapping. This resulted in a set of 15,139,199 SNPs. For each trait, we removed any SNPs with close to zero heritability using https://github.com/omerwe/polyfun/blob/master/munge_polyfun_sumstats.py.

We used two annotation sets to perform functionally informed fine mapping. The PolyFun-Baseline fine mapping runs used the baseline-LF version 2.2.UKB (baseline-LF) annotation set, consisting of 187 annotations including coding and conservation features as well as overlap with promoters and enhancers [10]. The baseline-LF annotation set was downloaded from https://data.broadinstitute.org/alkesgroup/LDSCORE/baselineLF_v2.2.UKB.polyfun.tar.gz. A complete description of this annotation set is available via PolyFun [8]. Sniff fine mapping combines the baseline-LF annotations with 102 variant effect annotations derived using Borzoi.

Functionally informed fine mapping was performed as described in PolyFun [8]. To compute LD scores, we used summary LD information derived from 337,000 British-ancestry individuals within the UK Biobank, made available by Weissbrod et al. with the PolyFun publication [8]. The British-ancestry LD reference was downloaded from https://alkesgroup.broadinstitute.org/UKBB_LD. Given this LD reference, we computed LD scores using the PolyFun scripthttps://github.com/omerwe/polyfun/blob/master/compute_ldscores_from_ld.py. We then ran PolyFun in its non-parametric setting to compute variant prior causal probabilities using the UK Biobank LD information. To perform genome-wide benchmarks, we followed PolyFun’s strategy of fine mapping overlapping 3 Mb loci (overlapping with a 1 Mb stride). We aggregated fine mapping results across these windows using the PolyFun codebase https://github.com/omerwe/polyfun/blob/master/aggregate_finemapper_results.py. For each SNP, we estimated its PIP by using the 3 Mb window within which it was most central.

### Fine mapping evaluation

Credible set entropy was computed as entropy of the distribution of variant posterior inclusion probabilities (PIPs) within the 95% credible set. Prior to computing entropy, variant PIPs within each 95% credible set were normalized to sum to 1.

### Defining Sniff-prioritized variant sets

Sniff-prioritized variants were defined using two criteria: (1) they were not fine mapped by either SuSiE or PolyFun-Baseline (PIP *<* 0.1), and (2) they exhibited increased PIPs when fine mapped with Sniff compared to PolyFun-Baseline (ΔPIP ≥ 0.1, 0.25, or 0.5). For each Sniff-prioritized variant, we defined a set of control variants as variants that were included in the same 95% credible set as the prioritized variant, but were assigned low posterior inclusion probabilities across all three fine-mapping methods (SuSiE PIP *<* 0.1, PolyFun-Baseline PIP *<* 0.1, and Sniff PIP *<* 0.1). At ΔPIP=0.1, we identified 6,834 unique Sniff-prioritized variants and 50,444 control variants. Of these, 1,541 Sniff-prioritized variants and 7,575 control variants were assayed in the Siraj MPRA data. At ΔPIP=0.25, we identified 1,158 unique Sniff-prioritized variants (of which 324 were assayed in the Siraj MPRA data) and 7,255 control variants (of which 1,386 were assayed in the Siraj MPRA data). At ΔPIP=0.5, we identified 299 unique Sniff-prioritized variants (of which 82 were assayed in the Siraj MPRA data) and 553 unique control variants (of which 154 were assayed in the Siraj MPRA data).

Differentially prioritized variants were defined as variants whose relative prioritization changed (i.e., they were either promoted or demoted) when fine mapped with Sniff as compared to PolyFun- Baseline. Sniff-promoted variants were defined as variants that exhibited increased PIPs when fine mapped by Sniff compared to PolyFun-Baseline, and Sniff-demoted variants were defined as variants that exhibited decreased PIPs when fine mapped by Sniff compared to PolyFun-Baseline (ΔPIP > 0.1, 0.25, or 0.5). At ΔPIP=0.1, we identified 11,644 unique Sniff-promoted variants and 10,055 unique Sniff-demoted variants. Of these, 3,282 and 3,033 variants were assayed in the Siraj MPRA data respectively. At ΔPIP=0.25, we identified 3,160 unique Sniff-promoted variants (of which 1,124 were assayed in the Siraj MPRA data) and 2,579 Sniff-demoted variants (of which 903 were assayed in the Siraj MPRA data). At ΔPIP=0.5, we identified 859 unique Sniff-promoted variants (of which 312 were assayed in the Siraj MPRA data) and 566 Sniff-demoted variants (of which 191 were assayed in the Siraj MPRA data).

### MPRA benchmark

To assess the functional activity of differentially fine-mapped variants, we downloaded MPRA data from Siraj et al. (2024) [46]. The authors measured the functional activity of 304,278 variants (or 608,556 alleles). The variant set consists of trait-associated variants that lie in credible sets and control variants. Activity was assayed across five cell-types—K562, SK-N-SH, HepG2, A549, and HCT116. We used the reported *log*_2_ fold changes as measures of allele-specific variant activity. From this set, we removed variants for which allele-specific activity measurements were missing in >2 cell types. We computed allele-specific activity as the average of the Δ log_2_ fold changes across the five cell types. If a variant was missing in≤ 2 cell types, we computed its allele-specific activity as the average Δ log_2_ fold changes across cell types in which variant activity was measured.

### FinnGen Replication

We downloaded FinnGen release 12 data from https://finngen.gitbook.io/documentation/data-download [47]. The FinnGen consortium performed fine mapping in 3 Mb windows around significant lead variants using SuSiE [5] and FINEMAP [4]. Fine mapped results from this cohort were available for two traits overlapping our trait set (height and eczema). Aggregated across the 2 traits, we extracted 80 Sniff-prioritized variants (Sniff ΔPIP>0.5) and 116 control variants from the same credible sets. Of this set, 85% had MAFs>0.1% in FinnGen. For Sniff-prioritized and control UKB variants which did not have MAF>0.1% in FinnGen, we assigned them PIP=0 in the FinnGen data (i.e., they were treated as non-replicated, Fig. 3H). The fraction of variants with PIP>0.1 in FinnGen were considered replicated and reported. We repeated the same process for Sniff-prioritized variants defined at thresholds of ΔPIP>0.25 and >0.1. For additional benchmarking, we focused on the subset of variants that had MAFs>0.001 in the FinnGen cohort, which resulted in 57 Sniff-prioritized variants (Sniff ΔPIP>0.5) and 109 credible-set matched controls.

### Heritability enrichment analysis with stratified LD score regression (S-LDSC)

S-LDSC assumes that each annotation independently and additively contributes to per-SNP heritability. Given a set of annotations {1, 2, …, *C*}, S-LDSC estimates the contribution of each annotation to per-SNP heritability as follows:

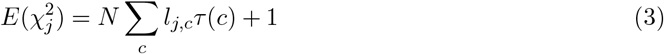

Here, *l*_*j,c*_ is the LD score of SNP *j* with respect to functional annotation 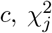 is the GWAS summary statistic for SNP *j*, and *N* is the GWAS sample size [42]. The estimated *τ* (*c*) values are interpreted as the contribution of an annotation to per-SNP heritability, or effect sizes of an annotation conditioned on all other annotations included in the regression.

For each of the 102 Borzoi PCs, we defined a set of 188 annotations, consisting of the 187 baseline annotations and 1 Borzoi PC. We performed S-LDSC given these annotation sets to estimate *τ* (*c*) values for each Borzoi PC conditioned on the set of 187 baseline annotations. We performed this analysis for all 15 traits derived from the UK Biobank, resulting in 102×15 *τ* (*c*) values, representing the conditional effect sizes of 102 Borzoi PCs on 15 traits. For the analyses in Figure 4, we focused on the estimated *τ* (*c*) values for the 13 Borzoi PCs derived from 87 GTEx RNA-seq VEPs, spread across 31 tissues. We used the S-LDSC implementation provided within the PolyFun package https://github.com/omerwe/polyfun/tree/master.

Having estimated the effect sizes, we computed standardized effect sizes *τ* ^*∗*^(*c*) as follows:

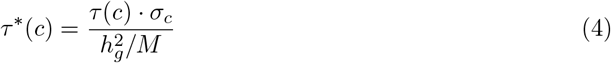

Here, *σ*_*c*_ represents the standard deviation of annotation 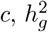 represents the total SNP heritability, and *M* is the number of SNPs used to estimate heritability 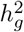. The standardized effect sizes *τ* ^*∗*^(*c*) therefore represent the proportionate change in per-SNP heritability given 1 unit standard deviation increase in the annotation value [36, 37].

### Gene assignment to credible sets

For each credible set *C*, we nominated a putative target gene as the gene with the highest proximity score to the credible set, weighted by variant PIPs. Let *V* = {*v*_1_, *v*_2_, …, *v*_*n*_} represent the set of variants in credible set *C*, where each variant *v*_*i*_ has genomic position *p*_*i*_ and PIP_*i*_.

For each protein-coding gene *g* on the same chromosome as credible set *C*, we computed a proximity score *S*(*g, C*) as follows:

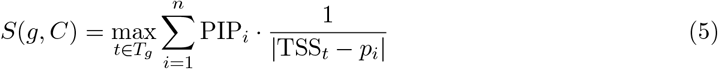

Here, *T*_*g*_ represents the set of all protein-coding transcripts for gene *g*, and TSS_*t*_ is the transcription start site position of transcript *t*.

The target gene *ĝ* for credible set *C* was then assigned as:

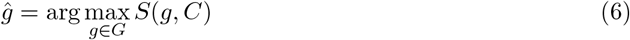

where *G* is the set of all protein-coding genes on the same chromosome as credible set *C*.

We note that in this formulation, for genes with multiple protein-coding transcripts, we computed a score *S* for each alternative TSS and selected the maximum score across all annotated transcripts, enabling us to nominate genes which might have an alternative TSS in close proximity to a credible set, even if the canonical TSS is distal to the credible set.

### Gene scoring with PoPS

We downloaded PoPS (v0.2) from GitHub: https://github.com/FinucaneLab/pops. PoPS relies on MAGMA to compute gene-level associations. We downloaded MAGMA (v1.10) from https://cncr.nl/research/magma and ran it using an LD panel derived from EUR ancestry individuals in Phase 3 of the 1000 Genomes Project, made available with the software. MAGMA requires gene annotations to compute gene-level associations. We downloaded the GENCODE v46 annotations transferred to GRCh37 (https://www.gencodegenes.org/human/release_46lift37.html). We subsetted the annotations to protein-coding genes with at least one protein-coding transcript, resulting in 19,839 genes. MAGMA requires a single transcript per gene. We chose to use the transcript that was annotated as the “MANE-selected” transcript. For genes that did not have a transcript with a MANE Selected tag (760 genes), we chose the longest transcript.

To run PoPS, we used the feature set provided in the paper, containing 57,742 gene features. We downloaded the feature set from https://github.com/FinucaneLab/gene_features. For the subset of genes that did not have features in this feature set, we set the PoPS “–nan policy” option to “mean”. This estimates missing feature values as the mean value of that feature across all genes. We ran PoPS using its default parameter settings except using “mean” for the “–nan policy” option. For each credible set, we nominated a gene as a PoPS target gene if it was the gene with the highest PoPS score within 1 Mb of the credible set midpoint. We calculated credible set midpoint as the mean position of all variants in the credible set.

## 6 Data and code availability

The pre-trained Borzoi model used to score variants was accessed via https://storage.googleapis.com/seqnn-share/borzoi/f0/model0_best.h5 [30]. Code used to score variants with Borzoi is available here: https://github.com/calico/baskerville/blob/main/src/baskerville/scripts/hound_snp.py.

We have released Borzoi variant effect predictions for a set of 19,534,182 common and low-frequency variants, which are available here: https://console.cloud.google.com/storage/browser/seqnn-share/sniff/borzoi_veps. The processed Borzoi-102 annotations for the set of 19.53 million variants are available here: https://storage.googleapis.com/seqnn-share/sniff/borzoi_102_annotation_set. All functionally informed fine mapping was performed using the PolyFun codebase: https://github.com/omerwe/polyfun [49]. Gene annotations (GENCODE v46) were obtained from https://www.gencodegenes.org.

## 7 Acknowledgments

This work was funded by Calico Life Sciences LLC. The funder had no role in study design, data collection, or analysis. Publication of the manuscript was approved after an internal scientific review process. We thank Madeleine Cule, Fanny Huang, Johannes Linder, David Wang, and Benjamin Auerbach for helpful discussions and valuable feedback.

## A Supplementary figures

**Figure S1:**
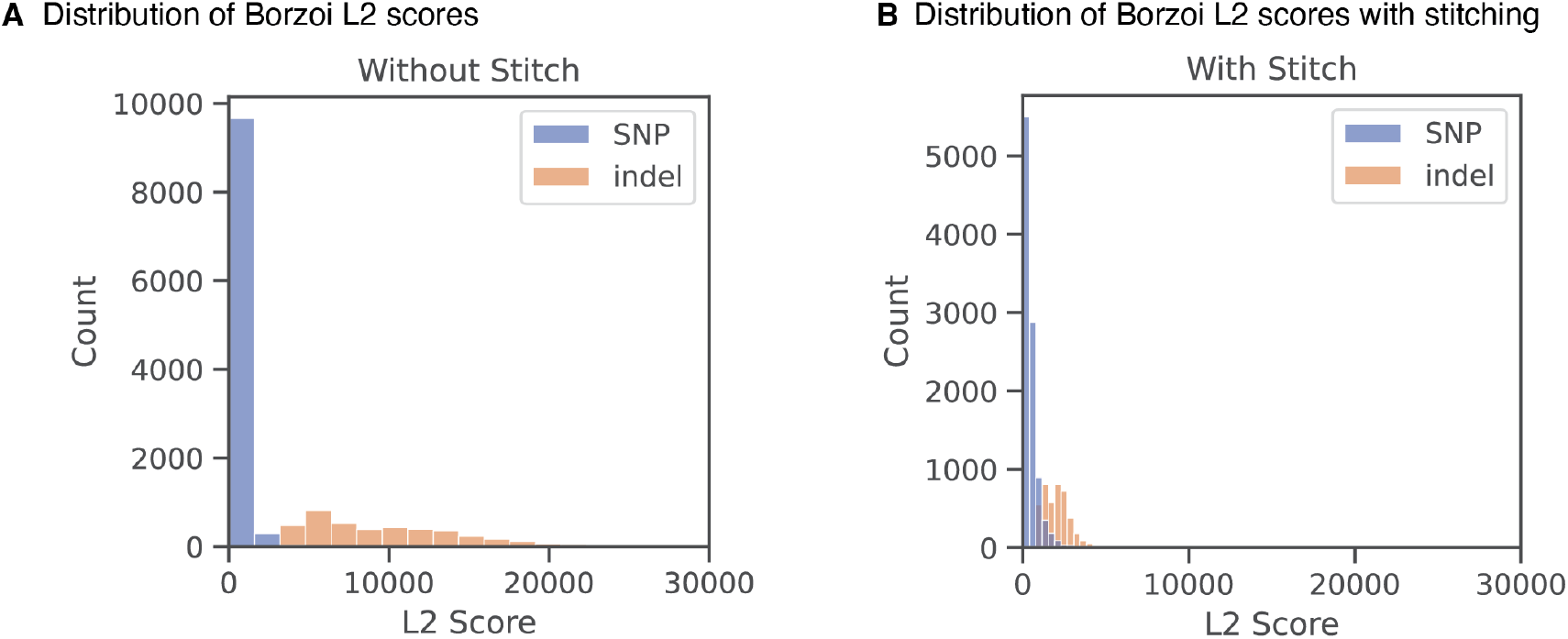
Comparison of Borzoi L2 scores for a subset of 50,000 sampled SNPs and indels with and without stitch-based indel scoring. L2 scores are summed across all output tracks. Stitch-based shift augmentations mitigate technical bias in the model’s indel L2 scores.

**Figure S2:**
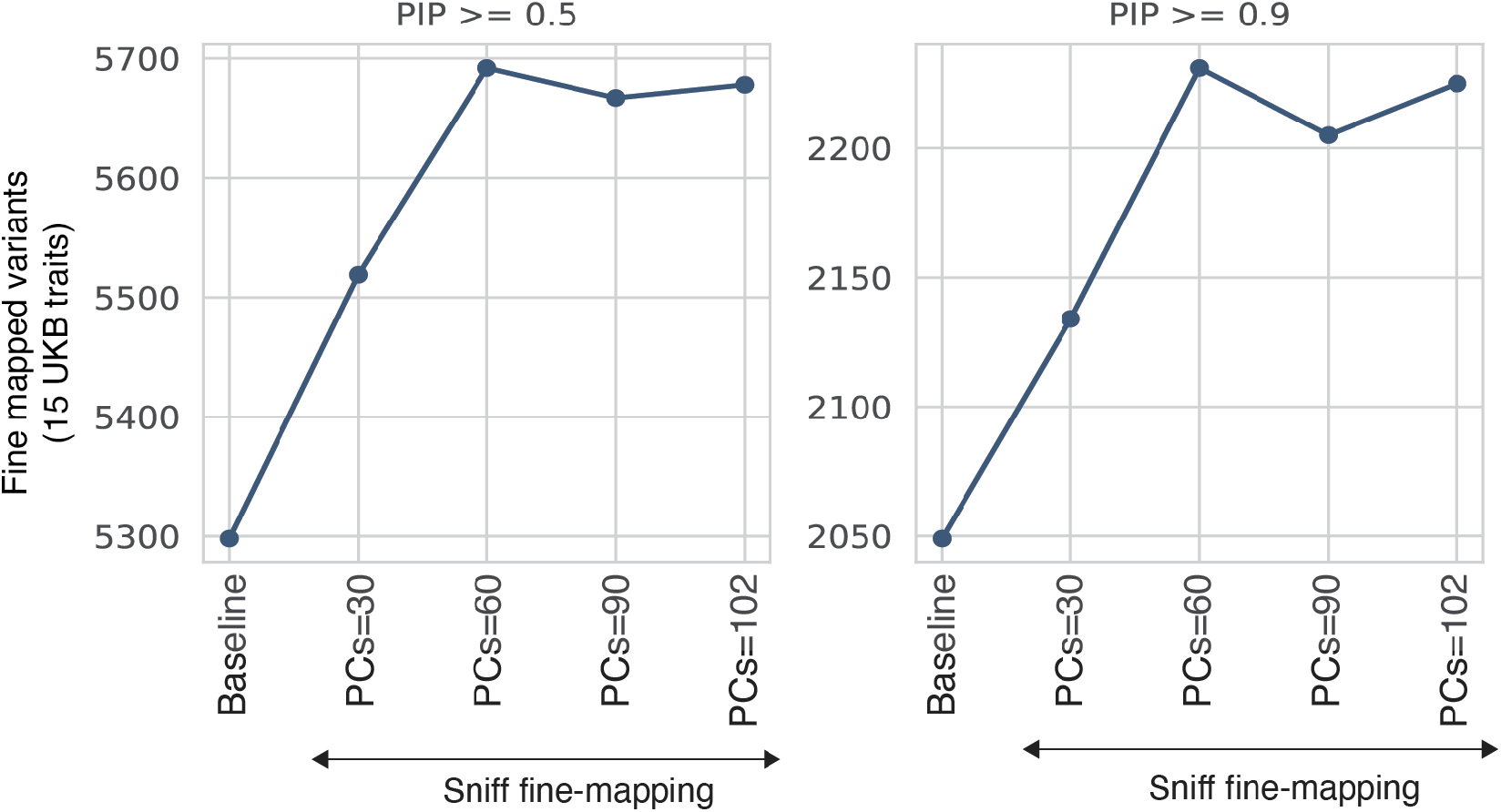
Number of fine-mapped variants, aggregated across 15 UKB traits. PCs=30 through PCs=102 refer to annotation sets derived by performing PCA per assay category and retaining 5, 10, 15, or the number of PCs that explain 95% of variance (for a maximum of 20) PCs per assay-category. Sniff fine mapping outperforms baseline-only fine mapping at PIP thresholds=0.5, 0.8 (Fig. 2A) and 0.9.

**Figure S3:**
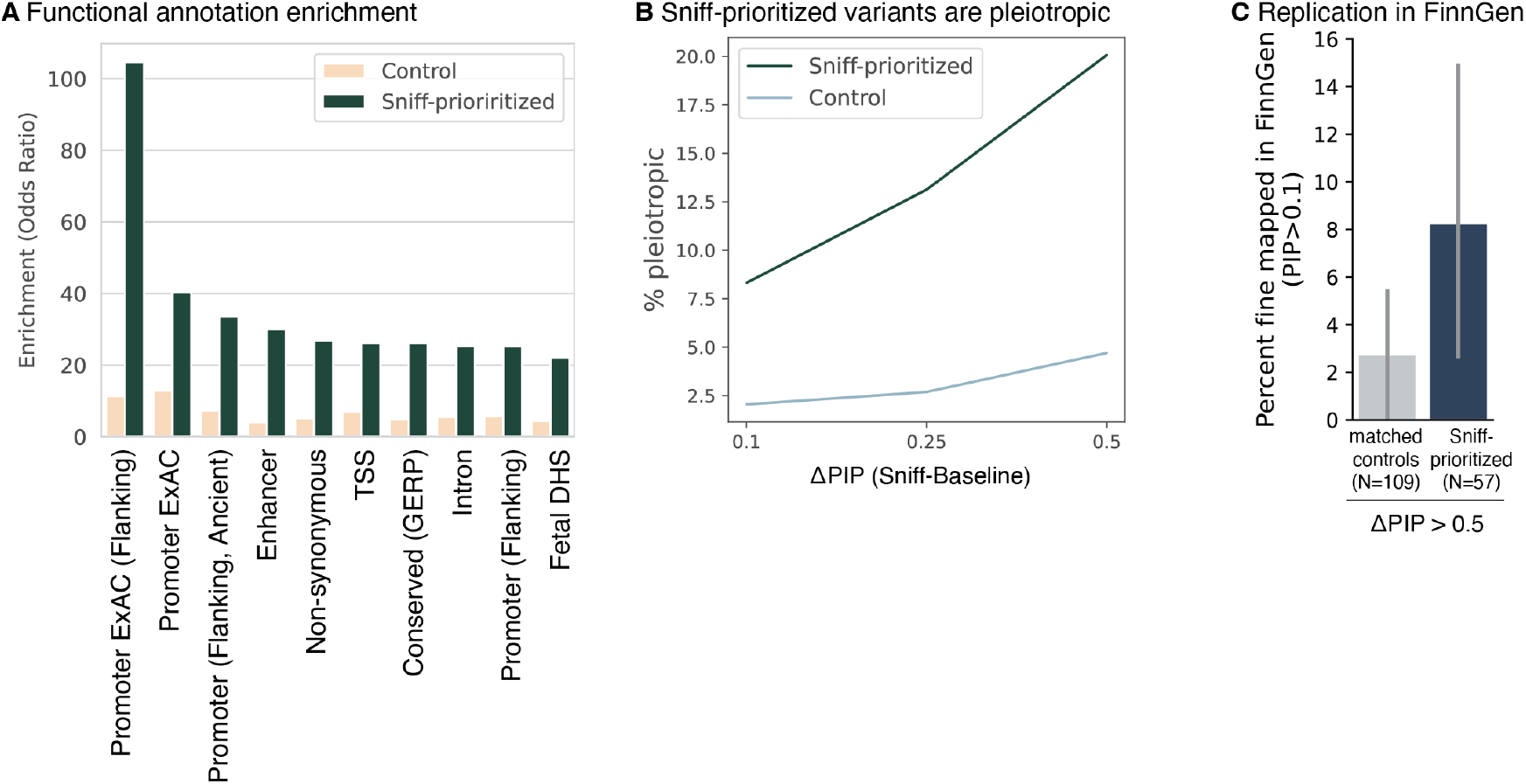
**A** Sniff-prioritized variants are significantly enriched (Fisher’s exact test *p <* 0.05) for baseline functional annotations including promoters, enhancers, non-synonymous nucleotides, transcription start sites (TSSs) and DNase-hypersensitive regions. In beige, enrichment is plotted for control variants from the same credible sets. **B** Percentage of Sniff-prioritized and credible set-matched control variants that are fine mapped in at least one other trait (among 94 fine mapped traits) with PIP>0.1.

**Figure S4:**
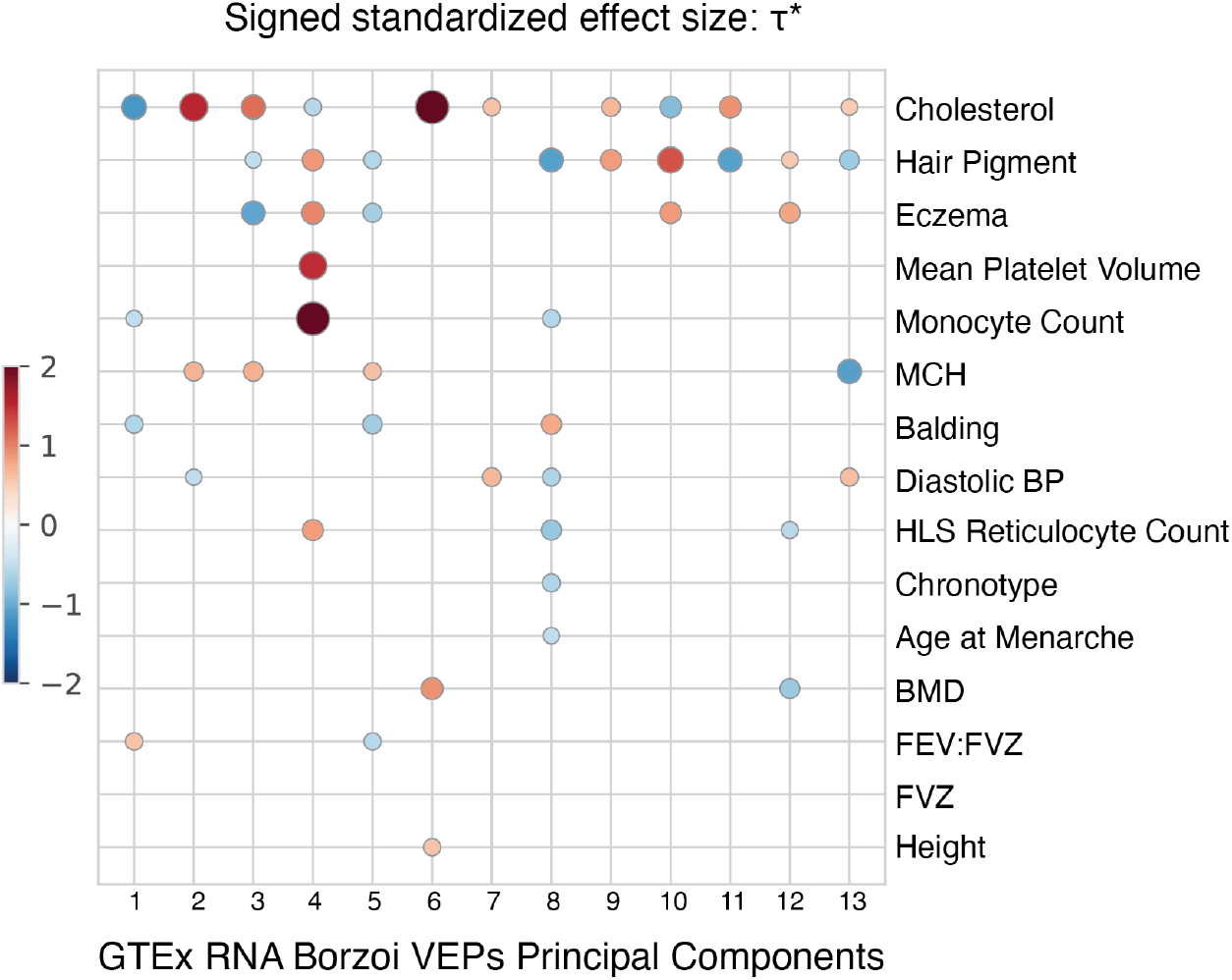
Signed standardized effect size *τ* ^*∗*^ for each Borzoi PC, computed by S-LDSC conditioned on 187 BaselineLF annotations. PC loadings are visualized in Figure 4A.

**Figure S5:**
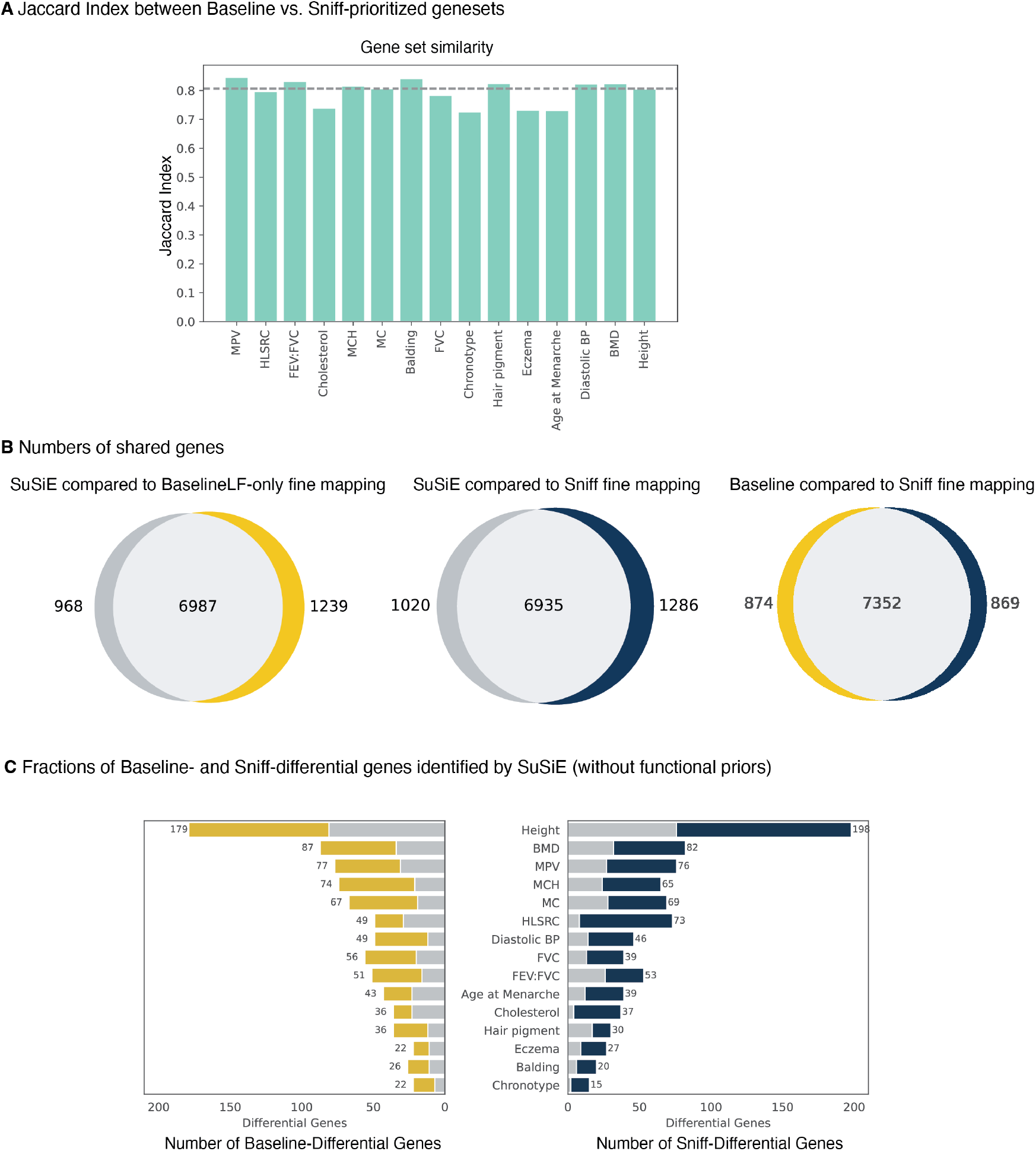
**A** Similarity between gene sets nominated by PolyFun-Baseline fine mapping and Sniff fine mapping as measured by the Jaccard index across the 15 UKB traits. **B** Venn diagram representing overlaps in gene sets identified by the three fine-mapping approaches: standard SuSiE (grey), PolyFun-Baseline (yellow) and Sniff (blue). **C** Bar plots represent number of genes that are differentially identified by PolyFun-Baseline and Sniff fine mapping. Gray bars represent the number of Baseline-differential and Sniff-differential genes that were also nominated by standard SuSiE.

## Notes

### Competing Interest Statement

D.S., A.K., Q.W., H.Y. and D.R.K. are employees of Calico Life Sciences. L.R. was an employee of Calico Life Sciences at the time this research was conducted.

